# An organotypic in vitro model of human papillomavirus-associated precancerous lesions allowing automated cell quantification for preclinical drug testing

**DOI:** 10.1101/2025.07.01.661539

**Authors:** Richard M. Köhler, Hans-Jürgen Stark, Iris Martin, Joel Altmann, Martin Simon Kalteis, Magnus von Knebel Doeberitz, Elena-Sophie Prigge

## Abstract

**Summary and graphical abstract:** 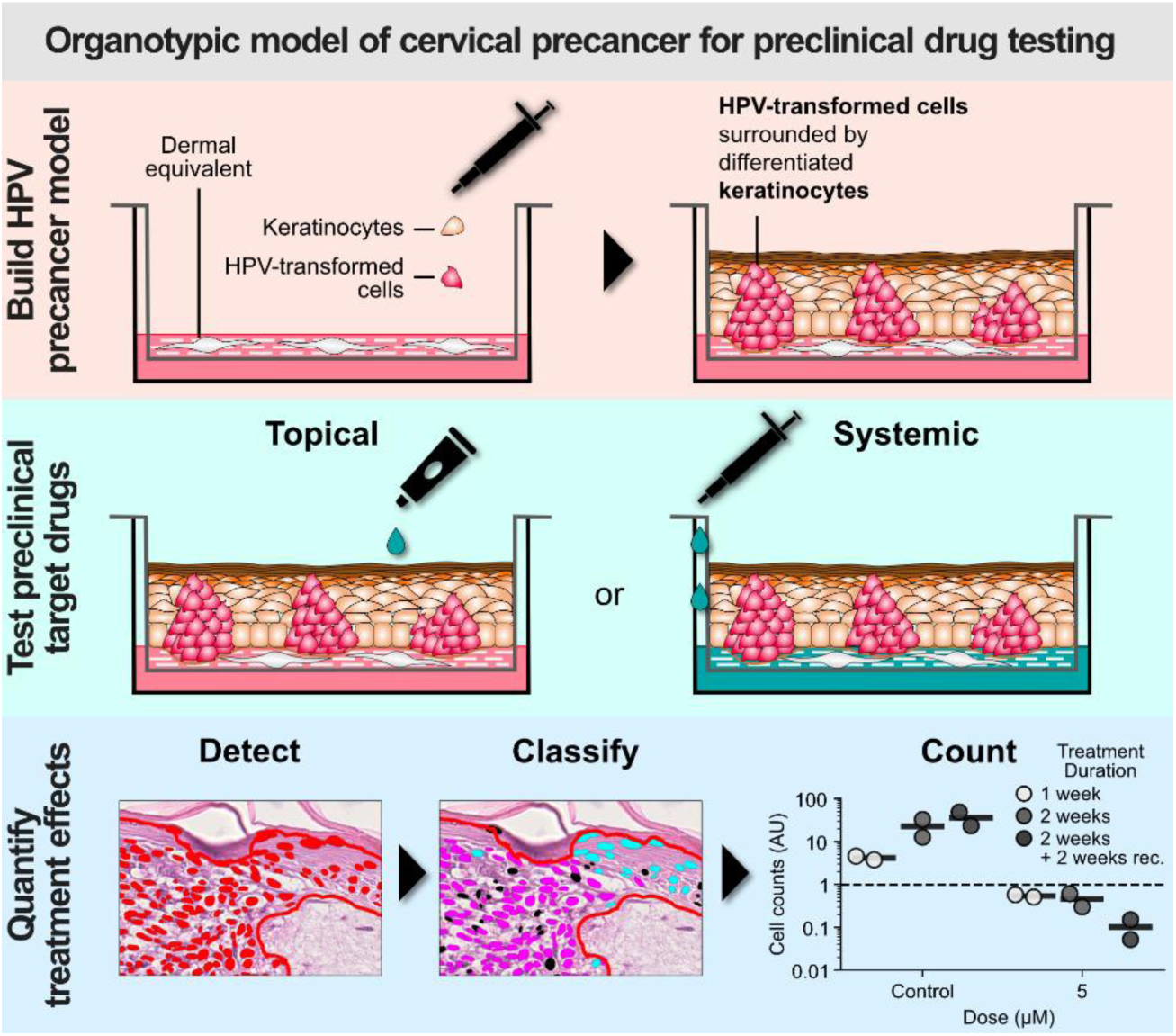

- A durable organotypic epithelial raft culture was established as a model of cervical precancer.
- Plausible time- and dose-dependent effects of cisplatin, 5-FU, and sinecatechins treatment were observed on keratinocytes and HPV-transformed cells.
- Treatment effects were reliably quantified using machine learning-based cell classification.
- This model may serve as a platform for preclinical investigation of topical and systemic treatment.

Oncogenic human papillomaviruses (HPV) are causally responsible for invasive cancers and precancerous lesions. These lesions represent a considerable disease burden worldwide, yet no causally effective treatments are available. The development of HPV tumor models realistically reflecting the *in vivo* treatment situation is necessary for finding new effective and tissue-sparing treatments. This study aimed to establish an *in vitro* model of HPV-induced precancerous lesions for preclinical drug testing and to provide an automated method for quantifying cell counts to assess treatment effects in this model.

To establish organotypic epithelial raft cultures (OTCs) as a model of HPV-induced precancerous lesions, we cultivated HPV-transformed cervical cancer cell lines SiHa, CaSki, HeLa, and SW756 in conjunction with primary human keratinocytes on a dermal equivalent and evaluated the impact of different cultivation variables. To demonstrate suitability of our model for preclinical drug application studies, we applied 5-fluorouracil, sinecatechins, and cisplatin either onto the air-exposed surface or to the growth medium, mimicking topical and systemic drug administration routes. We then developed a machine learning-based approach to quantify cell counts reflecting treatment effects in the established OTCs.

We successfully optimized a durable *in vitro* model of HPV-induced precancerous lesions. The model enabled monitoring of treatment effects in a three-dimensional context with differentiated consideration of the targeted HPV-transformed cells and surrounding normal epithelium. We demonstrated that extreme gradient boosted tree classifiers (XGBoost) can be successfully used to distinguish the number of tumor cells, normal keratinocytes, and degraded cells with high accuracy (81.8±8%; P=2×10^-6^). Critically, quantification results plausibly reflected microscopical observations and gave a fine-grained picture of time- and dose-dependent treatment effects.

The established model in combination with automated cell quantification can serve as a valuable tool in analyzing major preclinical endpoints in *in vitro* studies, such as target cell treatment efficacy, potential side effects, and treatment schedule optimization.

## Introduction

Oncogenic human papillomaviruses (HPV) account for 4.5% of the global cancer burden^1^ and are the causative agents in close to all carcinomas of the uterine cervix.^2^ As a result of the low coverage of prophylactic vaccination and insufficient cervical cancer screening programs in many less developed regions, the worldwide incidence of cervical cancer is expected to rise from 600,000 in 2020^3^ to about 1.3 million in 2069.^4^ The burden of HPV-associated precancerous lesions is even greater. Between 1995 and 2015, the prevalence of high-grade cervical intraepithelial neoplasia (CIN) was estimated at about 2% worldwide.^5^ Apart from CIN, HPV is associated with high-grade vulvar, vaginal, penile and anal intraepithelial neoplasia. In Europe alone, around 284,000 to 542,000 cases of anogenital precancerous lesions annually were estimated to be caused by HPV.^6^

The standard approaches for treatment of HPV-associated precancerous lesions consist of excisional or ablative methods using surgical techniques or laser,^7^ involving a significant risk for side effects.^8^ Despite considerable advances in the understanding of molecular processes during HPV-induced carcinogenesis, no causally effective therapy has been approved for these lesions to date. One factor impeding the development of targeted treatment options might be the paucity of suitable *in vitro* models of HPV-associated lesions to investigate the efficacy of novel therapies in a realistic setting.

While the complexity of tumor architecture and microenvironment is not adequately represented in conventional monolayer cell culture, animal experiments are resource-demanding and have been the subject of ethical and social debates aiming to limit their application in research. Consequently, more complex and representative *in vitro* models of HPV-related disease, such as three-dimensional (3D) culture systems, have been developed.^9^

As a model for precancerous lesions and invasive cancer, HPV-transformed cells have been seeded and cultured in organotypic epithelial raft cultures (OTCs) in the past.^10^ However, the interaction of different cell populations, such as keratinocytes and tumor cells, and observation of side effects – reflected by the toxicity of applied compounds on healthy keratinocytes – are of crucial importance. Yet we found but a single mention of the combined culture of HPV-transformed cells and healthy keratinocytes in an OTC context.^11^ Furthermore, conventionally used collagen gel matrices have a limited lifespan and limited physical stability that may impede the investigation of long-duration treatment regimens and topical application of creams or ointments.

In preclinical drug development, it is of great significance to obtain fine-grained estimates for the target drug dosage and required duration of planned treatment regimens. The study of immunostained sections of OTCs can provide a detailed picture of distinct treatment effects. However, it may be desirable to develop additional methods that provide quantitative measures of treatment effects to complement the qualitative assessment. A unique opportunity is now provided by the growing availability of powerful hardware and freely available software, which has accumulated in the advent of machine learning in various fields of natural science. We hypothesized that readily available hematoxylin and eosin staining combined with state-of-the-art machine learning models (XGBoost) could provide a reliable tool for automatized cell counting that may serve as a quantitative measure of treatment effects.

We aimed at creating a 3D *in vitro* model tailored to the investigation of novel treatment approaches for HPV-induced precancerous lesions in a preclinical setting, accelerating translation of findings to the human *in vivo* situation. Such a model should enable the quantification of short- and long-term effects of both systemic and topical treatment on transformed and normal epithelial cells in a 3D tissue architecture.

## Results

### Organotypic cultures morphologically reflect HPV-induced high-grade precancerous lesions after 2 weeks of culture

We aimed to create organotypic epithelial raft cultures (OTCs) mimicking HPV-induced precancerous lesions (Figure 1A-D). We therefore co-seeded primary human keratinocytes with SiHa cells in varying ratios (0.1, 1, and 10% SiHa) onto a scaffold-reinforced dermal equivalent that incorporates primary human fibroblasts. OTCs were harvested in weekly intervals over the course of four weeks. One week after epithelial cell seeding, the OTCs appeared macroscopically white, and had a largely smooth surface on macroscopic inspection (Figure 1E). H&E-stained sections revealed that the primary keratinocytes had formed a stratifying epithelium of few layers at this stage of cultivation (Figure 1G-I). Small islets of tumor cells were identifiable, largely proportional to the ratio of SiHa cells seeded. However, tumor cells had not yet organized in clearly distinguishable formations resembling high-grade precancerous lesions. After two weeks, we observed an epithelium displaying all layers of a fully stratified epidermis, including a thin cornifying layer and readily discernible nests of tumor cells in cultures with proportions of 0.1 and 1% SiHa cells (Figure 1J-K). Thereafter, SiHa cells appeared to gain a considerable proliferative advantage over the primary keratinocytes. In cultures with higher proportions of SiHa cells (i.e. 1 and 10%), the tumor cells had penetrated the basal membrane and invaded the dermal stroma and had overgrown large parts of the normal epithelium three weeks post-seeding (Figure 1N-O). After four weeks, almost no vital primary keratinocytes remained identifiable. SiHa cells now covered almost the entirety of the dermal equivalent and had grown invasively in all cultures, irrespective of the proportion of SiHa cells seeded (Figure P-R). OTCs remained viable over at least six weeks post-seeding, the longest observation period of this study (see Figure S1 for the fine-grained temporal evolution of an OTC up to six weeks). At this point, the cultures had a yellow coloring and had slightly detached from the sides of the inserts on macroscopic inspection (Figure 1F). We determined two weeks of culture after seeding of the epithelial cells as the optimal pre-cultivation duration before treatment start or harvest of OTCs in subsequent experiments, since a differentiated epithelium with discernible nests of tumor cells had formed at this point of time. This OTC architecture thus largely resembled the intended morphology of a precancerous HPV-induced lesion.

**Figure 1:**
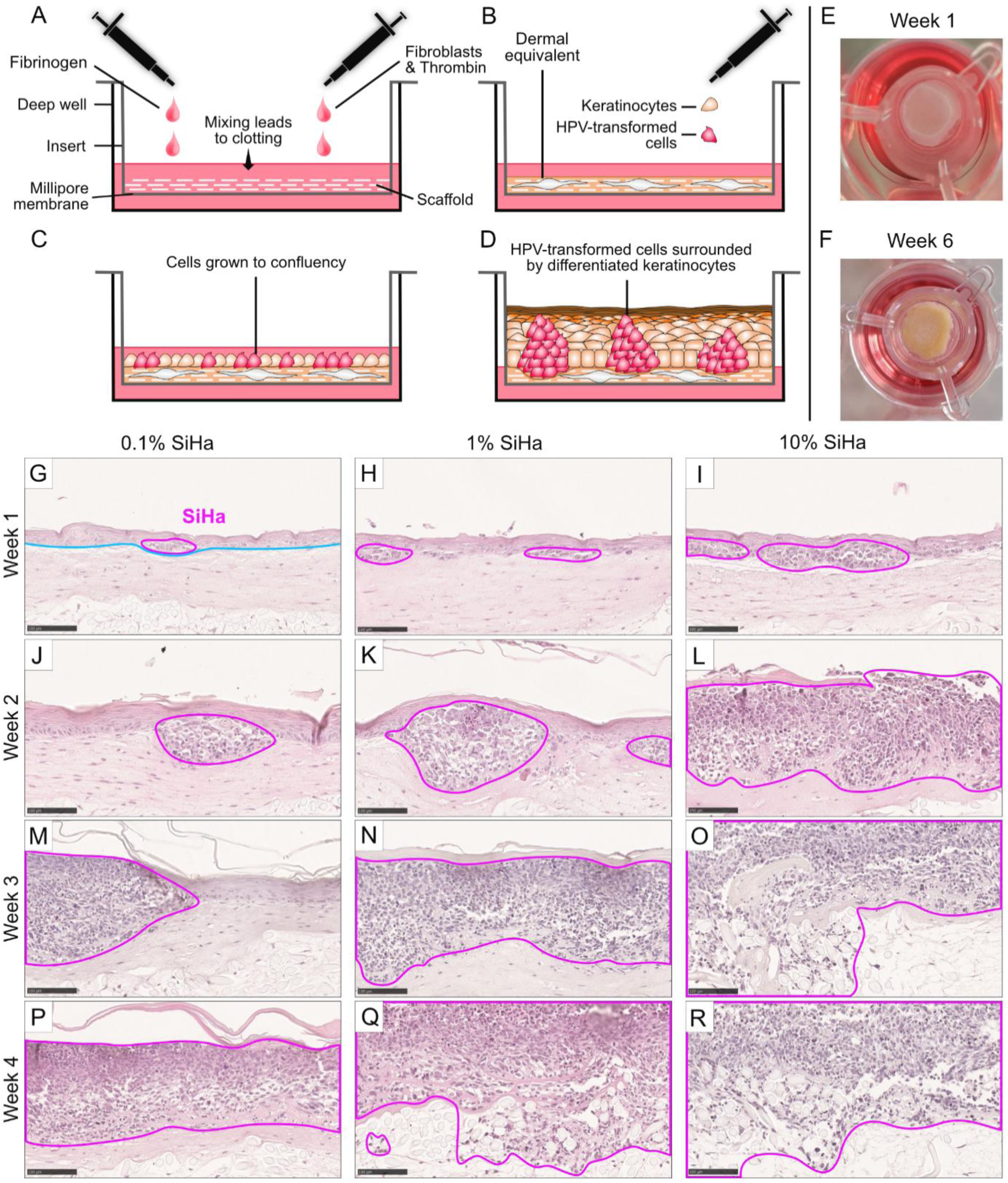
Establishment of organotypic epithelial raft cultures (OTCs) mimicking HPV-induced precancerous lesions. (A) Scaffold-enforced dermal equivalents were prepared by adding fibrinogen and a thrombin-fibroblast suspension onto a scaffold seated in inserts of a deep-well plate. (B) After several days, target keratinocyte−tumor cell mixtures were seeded onto the dermal equivalents. (C) After 24 hours of submerged culture, cells had grown to confluency, at which point the growth medium level was lowered to expose the OTC’s surface to air. (D) After about two weeks of culture, a fully differentiated epithelium had formed with discernible nests of tumor cells. (E, F) Macroscopic aspect (top view) of OTCs one and six weeks after seeding of epithelial cells on the dermal equivalent, respectively. (G – R) Images of H&E-stained sections of keratinocyte−SiHa co-cultures in different mixing ratios after one, two, three, and four weeks of culture. SiHa cells delineated in magenta. Dermal equivalent-epithelial border delineated in blue. Images depict representative tissue areas from one of three biological replicates per indicated treatment condition. Scale bars: 100 µm.

### Models of cervical precancerous lesions can be derived from various HPV-transformed cell lines

To account for the assumed biological and morphological variability of models deriving from HPV-transformed cells with different characteristics, we further established models using different HPV-transformed cell lines. Apart from HPV-16-transformed SiHa cells as described above, the models included HPV-16-transformed CaSki and HPV-18-transformed cell lines HeLa and SW756.^18–21^ Like SiHa cells, HeLa and SW756 cells grew in tumor formations easily distinguishable in H&E stained sections. In contrast, CaSki cells not only organized in tumor formations within the epithelium but also caused local erosion of the healthy epithelium (Figure S2). All HPV-transformed cell lines lacked morphological epithelial differentiation. However, a pronounced variability in growth rates was observed between different HPV-transformed cell lines and the ratio of seeded cells for the final models were adapted accordingly. HeLa cells expanded rapidly and a ratio of 0.1% tumor cells to normal keratinocytes was considered sufficient to generate an adequate model morphologically reflecting precancerous lesions after two weeks. In contrast, the cell lines SW756 and CaSki required a percentage of 5%, and SiHa a percentage of 1% tumor cells to achieve the desired morphological representation of HPV-induced precancerous lesions.

To facilitate the distinction of normal keratinocytes and HPV-transformed cells within OTCs, we characterized their nuclear and cytoplasmatic expression patterns of distinct target proteins. The combined detection of the cellular proteins p16^INK4a^ and Ki67 permits the highly sensitive and specific identification of HPV-transformed cells.^22^ In our model, p16^INK4a^/Ki67 co-staining allowed for a clear distinction between normal keratinocytes and HPV-transformed cell lines with only HPV-transformed cells staining positive for both markers (Figure 2). In normal keratinocytes, we observed a strong expression of keratin-14 in all layers except for the *stratum corneum*, whereas HPV-transformed cell lines showed a complete absence of keratin-14 (Figure 2). HPV-transformed cell lines HeLa, CaSki, and SW756 expressed keratin-7 and keratin-8. SiHa cells expressed keratin-7 but lacked keratin-8, while normal keratinocytes stained negative for both markers. Type IV collagen was chosen as a surrogate marker for the basal membrane and was strongly expressed below the basal layer of all specimens indicating the formation of a healthy basement membrane by primary keratinocytes (Figure 2). SiHa cell aggregates, however, led to a remarkable reduction of their underlying basement membrane when growing larger (Figure 2A).

**Figure 2:**
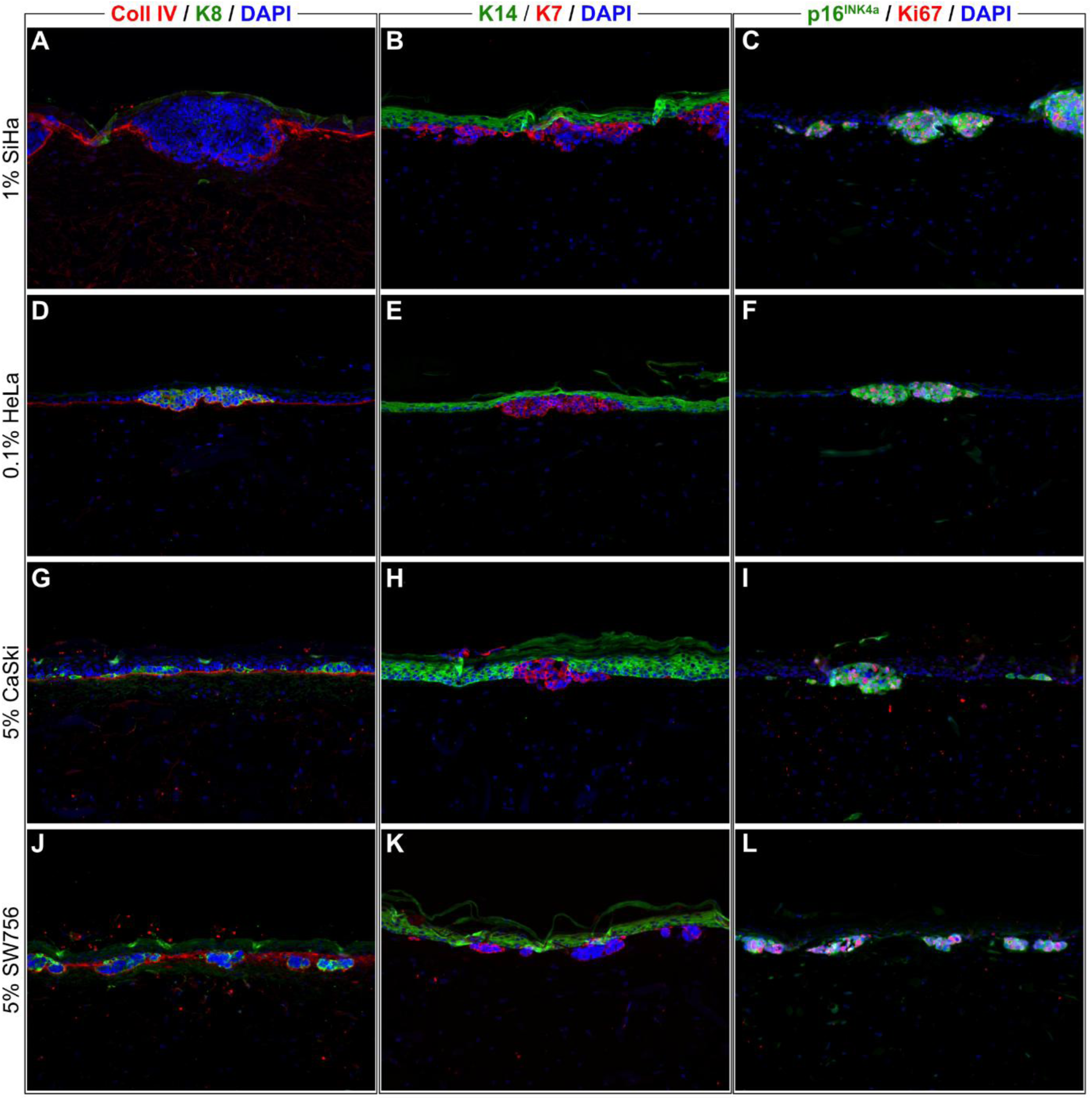
Phenotypic characterization of OTCs mimicking HPV-induced precancerous lesions with different HPV-transformed cervical cell lines. Immunofluorescent micrographs of OTCs derived from different cervical cancer cell lines after two weeks of culture (20x magnification). HPV-transformed tumor cells stained positive for keratin-8 (K8) – except for SiHa – and keratin-7 (K7), while keratinocytes stained positive for keratin-14 (K14). Collagen IV (Coll IV) was strongly expressed below the basal layer of the epithelium and indicated presence of a healthy basement membrane. HPV-transformed tumor cells additionally stained positive for p16^INK4a^ and Ki67, while normal keratinocytes did not. Images depict representative tissue areas from one of three biological replicates per indicated treatment condition. %: proportion of tumor cells seeded in relation to healthy keratinocytes.

### Time- and dose-dependent treatment effects can be observed in OTCs

We aimed to investigate the suitability of our models for preclinical studies investigating the effect of systemically or topically administered substances. To this end, we subjected OTCs to repeated treatments with either cisplatin applied into the medium, mimicking systemic drug administration, or commercially available 5-FU or sinecatechins ointments applied onto the air-exposed epithelial surface of the OTCs, both mimicking topical drug application (Figure 3). Before treatment, OTCs were kept in culture for about two weeks after epithelial cell seeding onto the dermal equivalent. OTCs treated with cisplatin were subjected to cisplatin added to the growth medium in different doses (1, 5, or 10 µM) for either one or two weeks, with a single treatment at the beginning of each week (Figure 3A). Additionally, a group of cisplatin-treated OTCs were kept in culture for 2 weeks after the treatment end (recovery phase). 5-FU or sinecatechins were applied to the air-exposed OTC surface for one and two weeks on five days per week (Figure 3A).

**Figure 3:**
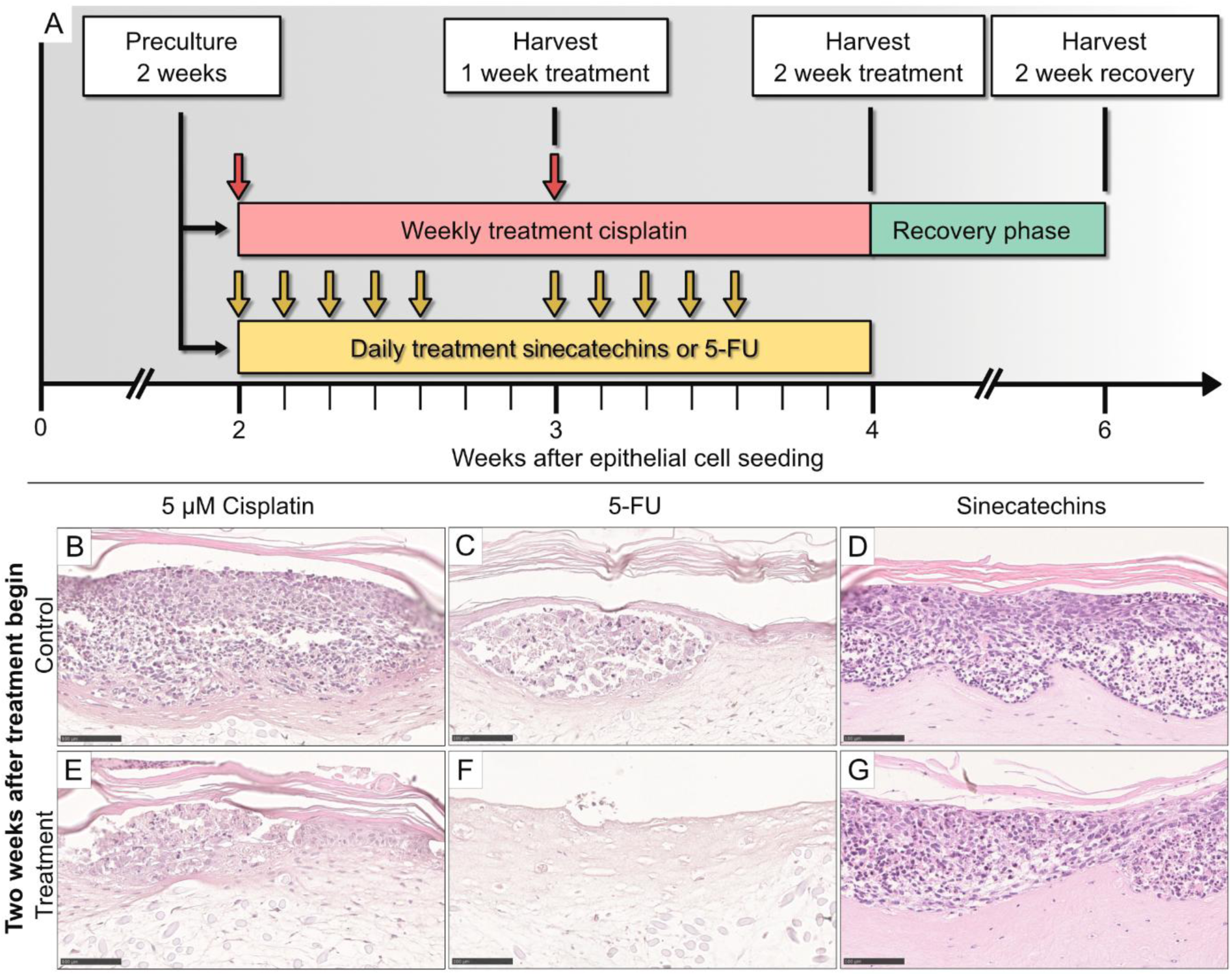
OTCs mimicking HPV-induced precancerous lesions as a model for testing topical and systemic treatment. (A) Before treatment begin, organotypic epithelial raft cultures (OTCs) were cultivated for about 2 weeks after co-seeding of 1% SiHa cells and 99% keratinocytes onto the dermal equivalent. OTCs were either treated with cisplatin added to the growth medium, with a commercially available 5-fluorouracil (5-FU) cream, or with a commercially available sinecatechins ointment applied to the OTCs’ air-exposed surface. Controls were left untreated. Cultures were harvested after one week of treatment, two weeks of treatment, or two weeks of treatment plus two weeks without further treatment (recovery phase). Example specimen after two weeks of treatment are depicted in panels B-G, see Figure S3 for the full panel of stained sections of all treatment durations. (B, C, D) In untreated controls, SiHa cells continued to proliferate. (E) Cisplatin addition to the growth medium led to degradation of SiHa cells compared to non-treated control cultures. (F) Daily administration of 5-FU cream caused complete detachment of the epithelium from the underlying dermal equivalent. (G) Topical application of sinecatechins led to a minor inhibition of SiHa proliferation and no detachment of the epithelium from the underlying dermal equivalent. Images depict representative tissue areas from one of 1 or 2 biological replicates per indicated treatment condition (see Methods for full details). Scale bars: 100 µM.

One week after the first application of cisplatin, a reduction in the number of SiHa cells could be observed in H&E-stained of all treatment groups (1, 5, and 10 µM) compared to untreated controls (Figure S3A). This effect was dose-dependent. The normal, stratifying epithelium remained morphologically unaffected by lower doses of cisplatin (1 µM and 5 µM). A reduction in the proportion of keratinocytes was observed in the group subjected to the highest dose (10 µM). One week after the second application of cisplatin, i.e. after two weeks of treatment, the extent of dysplastic formations had further reduced in treated cultures, whereas in untreated specimens, SiHa cells had largely overgrown the dermal equivalent and displaced the normal epithelium (Figure 3B, E). The observed cytotoxic effect of cisplatin on normal keratinocytes was more pronounced after the second week, with thinning of the epithelial layer occurring in all treatment groups. In the treatment group subjected to the highest dose of cisplatin, the effect was permanent, as the proportion of cells seemed to remain greatly reduced two weeks post treatment. Keratin-14 staining confirmed reduced proportion of SiHa cells (Figure 4A) and combined 5-ethynyl-2’-deoxyuridine (EdU) and cleaved caspase-3 staining revealed reduced cell proliferation and increased apoptosis in treated specimens (Figure 4 D).

**Figure 4:**
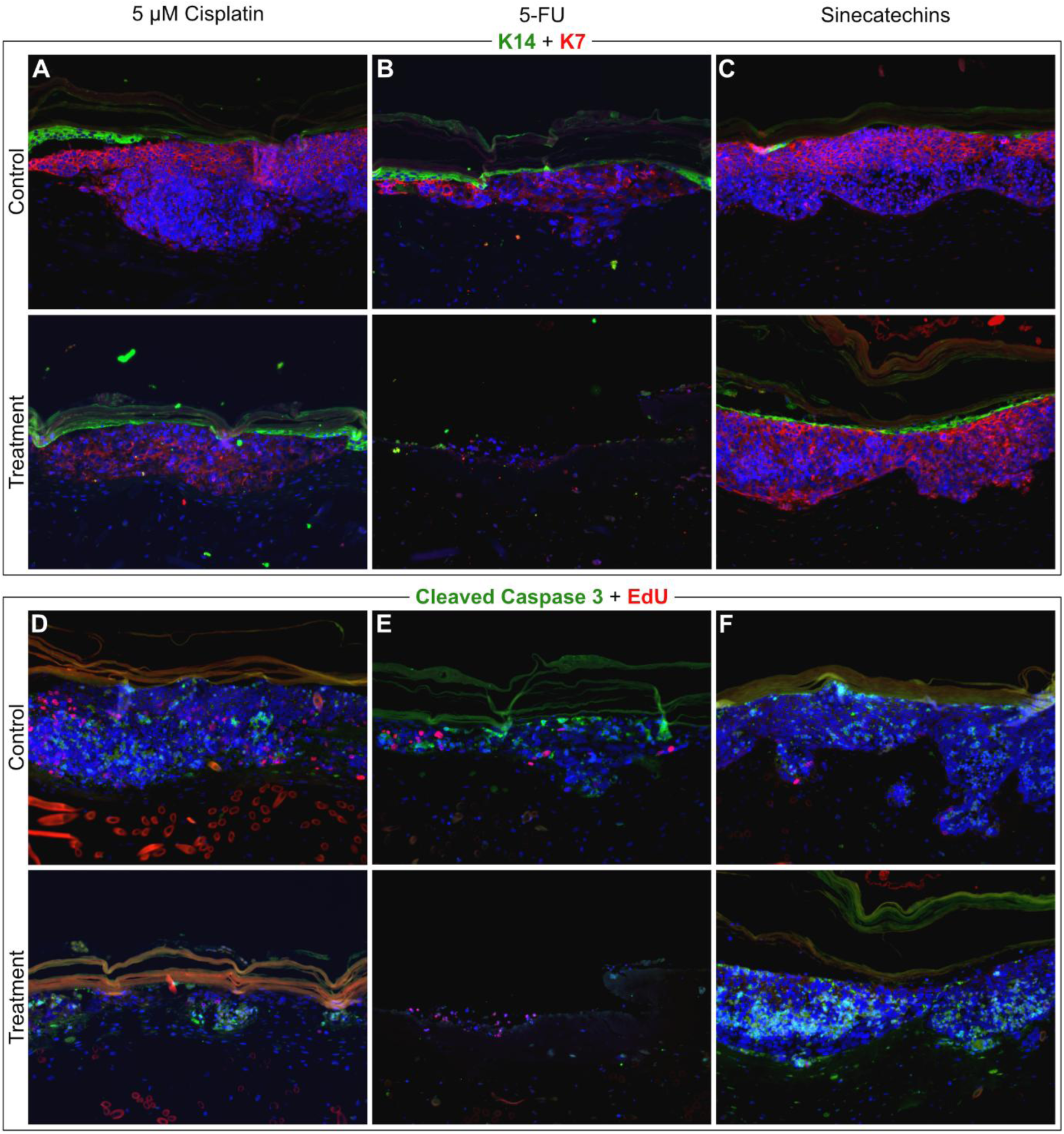
Differential treatment effects of cisplatin, 5-FU, and sinecatechins in OTCs mimicking HPV-induced precancerous lesions. Immunofluorescent micrographs of organotypic epithelial raft cultures (OTCs) at 20x magnification. OTCs were treated for two weeks with either cisplatin, 5-fluorouracil (5-FU), sinecatechins or left untreated as control. Keratin-7 (K7) and keratin-14 (K14) co-staining demonstrated degradation of SiHa cells in cisplatin (5 µM) and 5-FU treated cultures compared to untreated controls (A, B), while sinecatechins had only a minor effect on SiHa cells (C). Combined 5-ethynyl-2’-deoxyuridine (EdU) and cleaved caspase-3 staining demonstrated reduced cell proliferation and increased apoptosis after cisplatin treatment (D). Immunofluorescence staining reflects the detachment of epithelium with few remaining intact cells, as indicated by DAPI, after 5-FU treatment (E). A minor increase in apoptotic and a small decrease in proliferating SiHa cells was observed with sinecatechins treatment (F). Images depict representative tissue areas from a single of one or two biological replicates per indicated treatment condition (see Methods for full details).

### OTCs are suitable for topical drug application and reflect distinct treatment effects of different formulations

After one week of daily topical administration of 5-FU ointment, a reduction in cell number of both primary keratinocytes and SiHa cells was visible in H&E-stained sections, and the epithelium had detached from the dermal equivalent (Figure S3B). After a second week of daily 5-FU treatment, a near-complete loss of epithelial cells was observed, whereas in untreated controls, keratinocytes and SiHa cells were still present and tissue architecture was preserved (Figure 3C, F). This finding was corroborated by keratin-14 and keratin-7 staining, which revealed only few intact epithelial cells upon 5-FU treatment (Figure 4B). Nevertheless, EdU staining suggested that some actively proliferating SiHa cells remained (Figure 4E).

To investigate whether the detachment of epithelial cells after treatment was specific to 5-FU or the result of mechanical stress related to the application of a topical compound on the established OTCs, we subjected OTCs to daily treatment with a different topical compound, i.e. a sinecatechins ointment. Two weeks of treatment with sinecatechins ointment had only a small observable effect on the number of SiHa cells and keratinocytes in H&E-stained sections compared to untreated controls (Figure 3D, G) and the same observation was made in keratin-14/7-stained sections (Figure 4C). Cleaved caspase-3 and EdU staining suggested a slight increase in apoptotic and a minor decrease in proliferating SiHa cells (Figure 4F).

### Keratinocytes and tumor cells can be detected and reliably classified by a semi-automated approach in the newly established OTCs

To achieve a more fine-grained and objective representation of treatment effects, we aimed to quantify the number of viable cells in cultures treated with the different drugs and respective controls. First, we tested the validity of supervised classification of tumor cells and healthy keratinocytes in our model by automatically detecting objects in H&E-stained sections from the cisplatin treatment experiment (n=24 OTCs). These objects were then manually annotated as keratinocytes, SiHa, or degraded cells by two different authors. Then, XGBoost classifiers were trained to separate the three classes (keratinocytes, SiHa, degraded cells) from each other (Figure 5A). Detected objects were classified at an accuracy ranging from 66% to 97%, with a mean accuracy of 81.8±8% and 80.9±8% for independent annotations from two different authors (Figure 5B). The mean accuracy was therefore far above chance level (>33.3%; P=2×10^-6^), despite rigorous cross-validation. No significant difference in accuracy was measured for classifiers from the two researchers (mean Δ_Accuracy_ 0.9%; P=0.702).Visual inspection of individually classified objects revealed that the classifiers indeed assigned sensible labels to most detected objects (Figure 5C−H). High classification accuracy is therefore unlikely to be related to random assignment of correct labels by chance, suggesting the models may be used to derive a quantitative measure of treatment effects.

**Figure 5:**
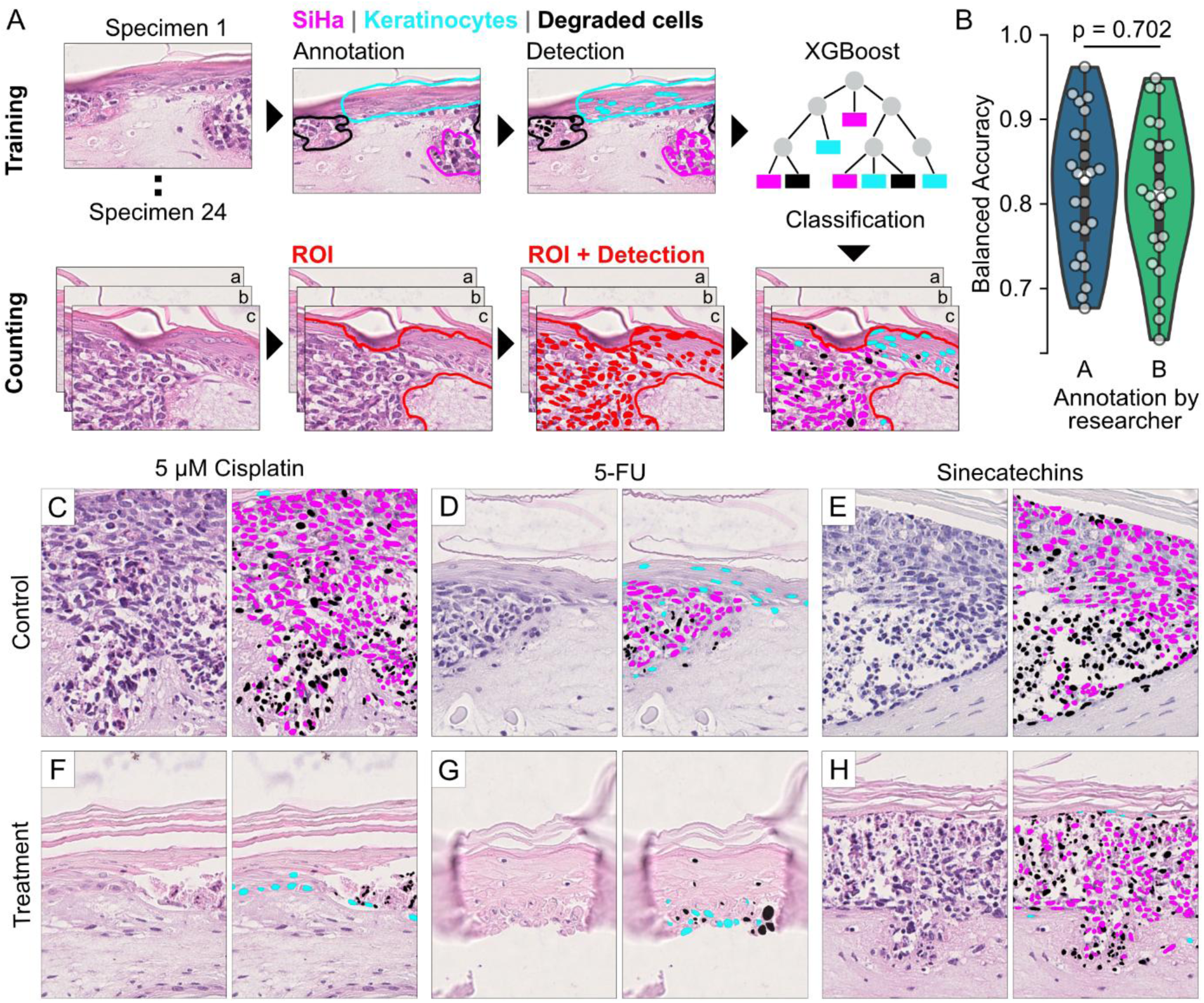
Quantification of treatment effects in OTCs through machine learning-based cell classification. (A) For classifier training, a single H&E-stained section was gathered from OTCs treated with cisplatin (N=24 specimens). After manual annotation of clusters of keratinocytes, SiHa cells, or degraded cells, nucleus-like objects (i.e., intact or fragmented nucleus) were automatically detected from star-convex polygon features using neural networks (StarDist) and used to train and validate XGBoost classifiers. During the counting step, these classifiers separated objects within the regions of interest (ROIs) of three additional sections of 100 µM distance from treated cultures from all treatment experiments (cisplatin, 5-FU, sinecatechins). (B) Separate annotations of cisplatin-treated specimen by two researchers A and B led to accuracies (mean ± std; N=24) of 81.8±8% and 80.9±8% (chance level 33.3%), without statistically significant difference during cross-validation (two-sided permutation test). A final classifier was then trained from all 24 specimen and applied to ROIs within specimen from all treatment experiments. Representative images without and with colored overlay of classified objects from untreated controls (C, D, E), and cultures treated for two weeks with cisplatin, 5-FU, or sinecatechins (F, G, H). Images depict representative tissue areas from a single of one or two biological replicates per indicated treatment condition (see Methods for full details). Colored overlay: magenta – SiHa; cyan – keratinocytes; black – degraded cells.

### Machine learning-based quantification reveals drug-, dose-, and time-dependent treatment effects

To derive a quantitative measure reflecting treatment effects, we investigated whether our classifiers could be used for semi-automated cell counting. A final classification model was trained using one section (of the total of four sections) per specimen from the cisplatin treatment experiment (24 in total), annotated by researcher A. The final model classified all detected objects in the annotated regions of interest in the remaining three sections per specimen. SiHa and keratinocyte cell counts in the treated specimens were normalized to the average of SiHa and keratinocyte cell counts in the untreated control specimens, respectively. Degraded cell counts were not considered, as debris from a single degraded cell can result in multiple object detections of varying counts and sizes, therefore not reflecting the actual number of degraded cells.

Cell counts reflected a clear time- and dose-dependent effect of cisplatin treatment in our OTCs. A single week of cisplatin treatment at the lowest dose (1 µM) reduced SiHa cell count to about 50% compared to untreated controls (Figure 6A). After two weeks of treatment and two weeks plus recovery phase, the cell count was reduced even further to around 35% and 30%, respectively. Higher doses led to an even stronger reduction in SiHa cells compared to 1 µM cisplatin but no pronounced difference was observed between the medium-dose and the high-dose conditions (5 and 10 µM). The ratio of SiHa to keratinocytes demonstrated that SiHa cells exponentially outgrew keratinocytes in untreated controls, indicated by values larger than 1, and that they continued to do so when treated with 1 µM cisplatin, although with a reduced growth rate (Figure 6G). However, when treated with 5 µM cisplatin, the *ratio* dropped below 1, and keratinocytes outnumbered SiHa cells. The relative growth advantage of keratinocytes further increased with longer treatment duration and after the two-week recovery phase (Figure 6G). By contrast, treatment with 10 µM cisplatin prevented the outgrowth of SiHa cells, but the effect was smaller than with 5 µM treatment, and no cumulative time-dependent effect was observed (Figure 6G).

**Figure 6:**
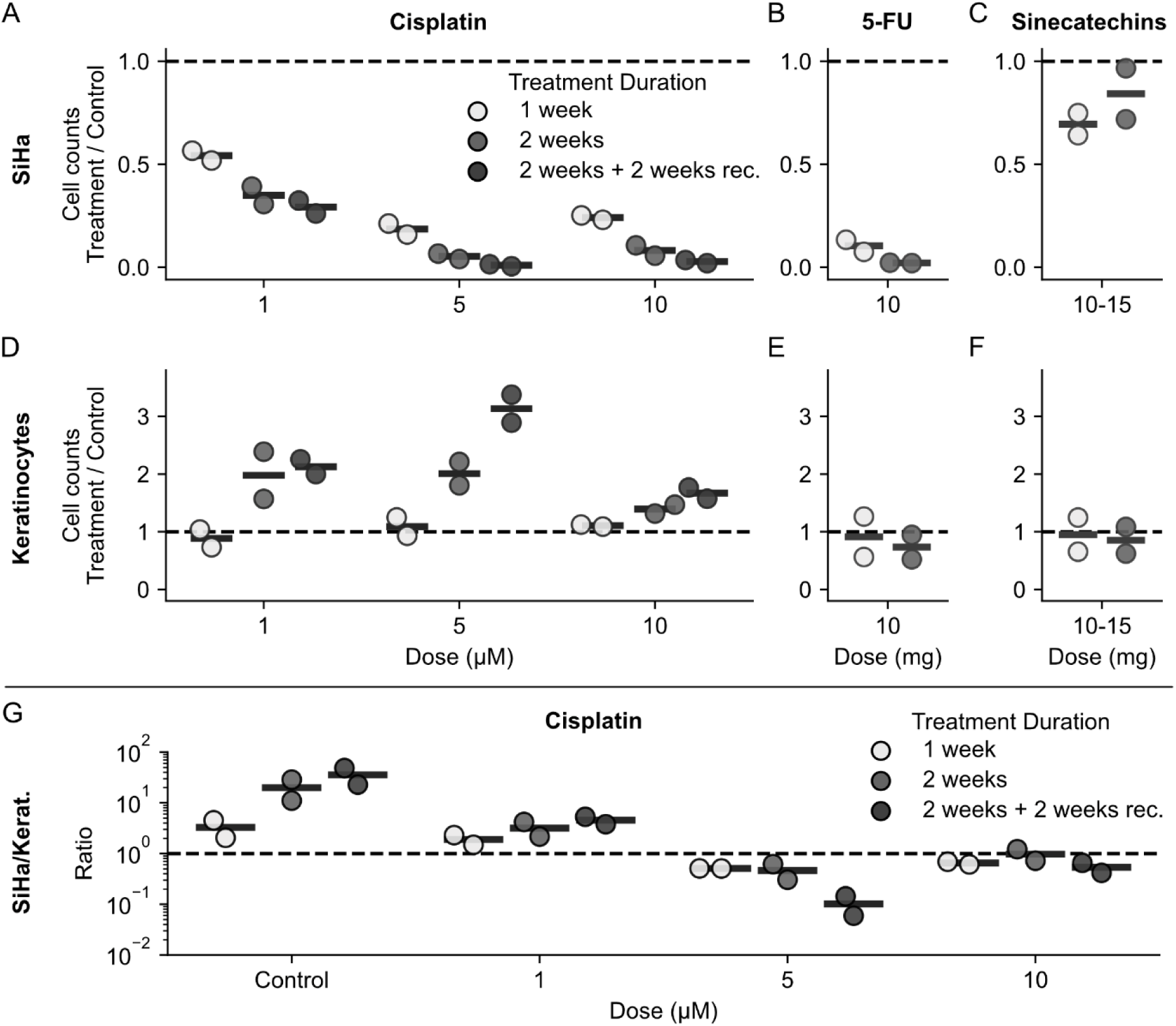
Machine learning-based quantification approach reveals drug-, dose-, and time-dependent treatment effects in OTCs when mimicking systemic and topical treatment. Panels show cell counts per mm^2^, normalized to untreated controls. Each dot represents the mean value of three sections from a single organotypic epithelial raft culture (OTC; two biological replicates per condition). (A-C) A reduction in SiHa cell count was observed for all treatment experiments. Longer treatment and recovery phase led to further reduction in SiHa counts, except for sinecatechins treatment. (D-F) A relative increase in keratinocyte count was observed after cisplatin treatment and was augmented by longer treatment and recovery phase. Only a small relative decrease in keratinocyte count was observed after two weeks of 5-fluorouracil (5-FU) and sinecatechins treatment. (G) The ratio of SiHa cells to keratinocytes demonstrated exponential outgrowth of SiHa cells in control specimens. After 1 µM cisplatin treatment, the exponential pattern persisted, albeit slightly reduced. 5 µM cisplatin treatment led to a growing proliferative advantage of keratinocytes. 10 µM treatment prevented the exponential advantage of SiHa cells but did not favor keratinocytes as strongly. Note that the y-axis in panel (G) is in logarithmic scale. No statistical tests were performed as the experiments were performed as proof-of-concept with low sample size (N=2 OTCs per condition).

Topical application of 5-FU for a single week caused a strong reduction in SiHa cells to less than 15% compared to untreated controls (Figure 6B). The two-week application schedule further reduced the number of SiHa cells. The number of primary keratinocytes after one week and two weeks of treatment was minimally lower than in controls (Figure 6E), although the epithelium had detached from the dermal equivalent after 5-FU treatment, so that these numbers must be interpreted with caution. In comparison to 5-FU treatment, the effect topical application of sinecatechins on SiHa cells was much less pronounced (Figure 6B). After a week of sinecatechins treatment, a reduction in SiHa count of about 30% and after two weeks of only 15% was observed. Sinecatechins application additionally resulted in a minimal reduction in keratinocytes relative to untreated controls (Figure 6F). In summary, we found that our quantification approach provided a fine-grained picture of treatment effects.

## Discussion

Recent analyses have predicted a rise in the burden of HPV-associated tumors in future years,^4^ while causally effective therapeutic approaches are still lacking.^7^ To bridge the gap from basic research and drug discovery to human translation, we aimed to establish a model for preclinical testing of novel treatment options that reflects high-grade precancerous lesions of the uterine cervix. We successfully created organotypic epithelial raft cultures (OTCs) comprising nests of HPV-transformed cells surrounded by healthy stratifying epithelium on a robust matrix containing vital fibroblasts, enabling the observation of multiple cell populations in a single model. The application of three different compounds, used for treatment of HPV-associated diseases (cisplatin, sinecatechins) or investigated in clinical studies (5-FU), added to the growth medium or onto the air-exposed surface, demonstrated that the model is suitable for the investigation of different treatment approaches. Additionally, we successfully established a semi-automated machine learning-based quantification approach that accurately reflected cell counts in our model. These findings suggest that the established OTCs have the potential to be used as a valuable model in preclinical drug discovery and research.

HPV-transformed cells in our established OTCs displayed a highly disorganized architectural phenotype showing no morphological epithelial differentiation, as has been previously observed in other raft cultures of HPV-transformed cell lines.^10,23,24^ However, our literature research yielded no previously published in-depth reports – only a single mention – on the combined culture of keratinocytes and HPV-transformed cells in organotypic raft cultures.^11^ The HPV-transformed cell lines HeLa, SiHa, and SW756 established dysplasia-like formations that were easily distinguishable from surrounding normal epithelium in our model upon a pre-cultivation period of about two weeks. However, in studies using this model, the quantity of viable HPV-transformed cells or keratinocytes might become greatly reduced by treatment and be limited to single cells or small cell clusters over time. The identification of the distinct cell populations may consequently prove to be more difficult. Likewise, after four weeks of culture in non-treated OTCs, SiHa cells had largely overgrown the surrounding epithelium leaving few remaining normal keratinocytes (irrespective of the proportion of SiHa cells at seeding), possibly impeding the analysis of treatment effects on normal keratinocytes. Therefore, to identify single cells and small cell clusters more accurately, we characterized the expression of different cellular proteins of HPV-transformed cell lines and primary keratinocytes in our OTCs. A combination of keratin-7 and keratin-14 staining permitted the reliable distinction of all our HPV-transformed cell lines from normal keratinocytes. As an alternative, keratin-8 proved to be a good marker for the used HPV-transformed cell lines with the exception of SiHa.

We evaluated the applicability of our model for preclinical research involving topically and systemically administered substances. OTCs were subjected to either cisplatin, a commercially available 5-fluorouracil (5-FU) cream (Efudix 5%) or a commercially available sinecatechins ointment (Veregen 10%). We observed the OTCs over the course of up to four weeks after the onset of treatment, adding up to a total culture duration of six weeks. Due to the long-term stability of our dermal equivalent, we were able to show that epithelial cells can remain viable for a considerable period in our model. Previous experiments making use of this dermal equivalent have shown that it supports even longer culture duration of twelve weeks.^25^ This aspect will prove to be useful when testing parameters relevant for future *in vivo* treatment, as treatment over a prolonged period of time is common in systemic as well as topical treatment regimens. Our model may be adopted into any laboratory’s workflow and holds potential in preclinical drug testing alongside animal models and monolayer culture. Animal models have advantages, such as a high degree of complexity, while OTCs have merits in lower financial and time expenses than animal models, but higher complexity than monolayer culture. Each model may therefore serve its purpose as a different link in the drug development chain.

Cisplatin is a chemotherapeutic drug that is applied as a key compound in the standard treatment of advanced and recurrent disease of cervical cancer.^26,27^ To simulate systemic treatment, cisplatin was added to the growth medium of our OTCs. It inhibited cell proliferation and induced cell death of SiHa cells and of primary keratinocytes, indicating that it was successfully able to penetrate the dermal equivalent and affect the different cell populations within the epithelial layer. The observations in our model were comparable to those previously reported for monolayer culture studies, in which treatment of HPV-transformed cells with cisplatin led to reduction in cell viability and induced apoptosis.^28,29^ Likewise, effects were similar to those observed after cisplatin treatment *in vivo*,^29,30^ indicating that our model may be useful in predicting the *in vivo* effectiveness of systemically applied drugs. The time- and dose-dependent effect of our results suggests that the model may be useful in treatment dose- and duration-finding experiments.

Currently, causally effective treatments for HPV-induced neoplasia of the cervix are lacking, while the number of cervical cancer cases worldwide have been estimated to double between 2020 and 2069.^4^ The treatment standard for HPV-induced precancerous lesions consists of excisional and ablative approaches,^7^ which involve a considerable risk for side effects.^8^ The risk is further aggravated by repeated interventions due to recurrent lesions. For these lesions, the development of topical formulations containing causally effective compounds would likely provide great benefits, serving as an alternative therapeutic approach. But despite efforts involving various compounds that have been tested in small clinical trials,^31^ no topical treatment has been approved to date. Imiquimod, that has demonstrated some efficacy for the treatment of high-grade cervical intraepithelial neoplasia (CIN 2/3) and vulvar epithelial neoplasia,^32^ is occasionally used as an off-label treatment for HPV-associated precancerous lesions. To validate our model for suitability of topical application, we subjected OTCs to treatment with either a 5-FU cream or sinecatechins ointment. 5-FU is an antimetabolite drug that causes DNA damage, leading to reduced DNA replication and cell death. 5-FU creams are currently used in the treatment of various dermatologic conditions such as actinic keratosis, Bowen’s disease, superficial basal-cell carcinoma, and warts.^33^ It has shown effectiveness in a preliminary clinical trial for treatment of CIN 2, with no moderate or severe side effects being reported in the 29 treated patients.^34^ However, others reported severe ulcerations following 5-FU cream application for the treatment of HPV-associated vaginal lesions.^35^ This may be related to the fact that whereas the former study reported outcomes for regimens with applications every two weeks, the latter employed daily and weekly application regimens with much higher total doses of 5-FU. The amount of side effects is thus likely to be dependent on the frequency and dose of administration. Similarly, after daily treatment in our experiments, we observed detachment of the epithelium, reflecting the effect reported with daily application *in vivo*.

Veregen is a topically applied ointment that has been approved for the treatment of benign HPV-associated genital warts.^36^ Its main agent is sinecatechins, a drug substance derived from an extract of green tea leaves which contains among other polyphenols the component epigallocatechin gallate (EGCG). Several studies have suggested EGCG as the main active agent in sinecatechins, and have demonstrated the antineoplastic activity of EGCG in HPV-transformed cervical cancer cell lines.^37^ It was shown to downregulate E6 and E7 expression in CaSki and HeLa cells,^38^ and there is evidence that its antineoplastic activity may be mediated through demethylation via inhibition of DNA methyltransferases.^39^ The safety of sinecatechins ointment for the use in usual type vulvar intraepithelial neoplasia (uVIN) has been demonstrated in a phase II study, where it did not lead to histological resolution, but induced a significant clinical response.^40^ In the current study, we observed that both sinecatechins and 5-FU were able to penetrate the *stratum corneum* of our OTCs, affecting both tumor cells and normal keratinocytes. Critically, our results demonstrate that the OTCs can withstand significant physical strain during topical application and subsequent removal of residues. The model may therefore serve as a valuable predictor of a compound’s ability or a specific preparation to penetrate the epithelium. It may also be used to compare different therapeutic strategies, such as testing different formulations of a single topical treatment.

The primary keratinocytes used in our model stratify into a keratinizing squamous epithelium as found in the epidermis of the human skin. Keratinizing epithelium differs in various aspects from the columnar or non-keratinizing squamous epithelium of the human cervix. The *stratum corneum* offers a barrier against not only mechanical but also chemical stress from the environment. The penetration of therapeutic compounds into the epithelium may therefore differ between keratinizing and non-keratinizing epithelia and the effect observed in experiments simulating topical application might not be readily transferable to the cervical non-keratinizing epithelium. However, we observed that the squamous layer only started to develop during the second week post-seeding, and as treatment in our experiments was initiated between day 11 and 13 post-seeding, the squamous layer was not strongly developed at this point. An alternative to epidermal keratinocytes may be the use of human cervical keratinocytes. However, the primary culture of normal cervical keratinocytes is not widely established due to low yields in cell numbers and frequent contamination with fibroblasts.^41,42^ Although some commercially available primary cervical keratinocytes exist, they are thus not as readily available as epidermal keratinocytes (e.g., from human foreskin tissue) and we opted for the use of primary epidermal keratinocytes. In all primary keratinocytes, the growth rate variability between strains from different individuals is non-negligent. Consequently, OTCs making use of primary keratinocytes will also display a certain biological variability, as relative proliferation rates compared to HPV-transformed cells and thus the ideal proportion of keratinocytes to HPV-transformed cells may vary among different batches. Culture parameters, such as pre-cultivation period and epithelial cell seeding ratio, should therefore be optimized, or at least verified for each batch of primary keratinocytes. But the variability among different keratinocyte batches may also be leveraged to ensure that treatment effects are consistent and replicable across keratinocytes derived from different human subjects. More uniform models may be created from immortalized keratinocyte cell lines such as HaCaT,^43^ which have more consistent proliferation rates. As a major drawback of this approach, *in vitro* epithelium formed by immortalized keratinocytes as well as their growth rates are less similar to that of epithelium *in vivo* than primary keratinocytes. It is important to note that while our model shows great similarities to human *in vivo* epithelium, there are obviously several differences. For example, during the application of topical compounds to the OTC surface, one must proceed with great care since the model’s epithelium is more sensitive to manual manipulation than normal human epidermis. The more viscous the topical formulation, the greater the risk that its application could lead to friction and consequent detachment of the epithelium from the dermal equivalent Therefore, we first applied the topical formulations onto a sterilized carrier film before administering it to the OTCs. Nevertheless, one must keep in mind that the repeated application of topical compounds as well as culture durations over several weeks may also increase the risk of microbial contamination of cultures.

To reliably demonstrate a compound’s effectiveness, it may be desirable to generate large numbers of OTCs in order to obtain adequate sample sizes. For the evaluation of treatment efficacy, a method for quantification of effects that can be readily applied to a large number of samples would thus provide a great benefit. Machine learning-based practical applications have recently been multiplying and have the potential to support humans in repetitive and laborious tasks – but also to aid in the field of medicine, for example in the assisted diagnosis of cancer from cervical imaging.^44^ Our findings demonstrate that it is sufficient to annotate regions in a small number of H&E-stained sections to train a model that could apply the learned patterns to other samples. It was able to accurately distinguish SiHa cells, keratinocytes, and degraded cell material. To generate SiHa and keratinocyte cell counts in other specimens, only the manual annotation of regions comprising cells of epithelial origin was necessary. Recent studies suggest that even the step of identifying target regions could potentially be automated through deep-learning based segmentation models.^45,46^ In this study, we accurately classified and discarded objects that were identified as degraded cell material. We did not analyze the degraded cell material counts, as a single degraded cell may result in multiple fragments and thus multiple objects of degraded material, and we did not feel confident that this number would yield a representative measure of degraded cells. Their detection was nonetheless necessary in order to avoid false additional cell counts for keratinocytes or SiHa cells. It is noteworthy that our results were obtained on a standard PC (4-core CPU, 32GB RAM, no dedicated GPU) and can therefore be translated to any laboratory without access to high-performance computing. Further, in this study we only made use of standard H&E-stained sections which are inexpensive. It is highly likely that slightly more expensive and time-consuming immunostaining in combination with deep learning would yield both even better detection of cell nuclei,^47^ and yield even higher classification accuracies for example using cell-type specific markers discussed in this study, such as co-staining of p16^INK4a^ and Ki67 or keratin-7 and keratin-14.

Finally, our OTCs are not restricted to the cell types presented in this study. For example, the incorporation of immune cells into organotypic cultures has already been demonstrated.^48,49^ The differential observation of local immune response may be desirable as immune evasion and immunosuppression are important factors in HPV-associated carcinogenesis.^50^ Modulation of the immune response for treatment of HPV-associated disease, for example via adoptive cell therapy or therapeutic vaccination, is an active field of research with ongoing clinical trials.^51^ It is important to note that the morphologic similarity of our cultures with precancerous lesions of the human cervix does not automatically implicate functional relevance. For example, the used cancer cell lines have been cultivated for many years and are likely to differ substantially from dysplastic cells *in vivo* with regards to proliferation rate, genome, metabolism, etc. On the other hand, their tumorigenic potential *in vivo* has been confirmed, for example through transplantation experiments into immune incompetent mice. Our model, as any model, does not claim to be a perfect replication of the *in vivo* situation, but we argue that it is an approximation that can be useful in many research situations. Therefore, in combination with our approach of semi-automated cell quantification, it holds the potential to be adopted by other laboratories in their research toward new therapeutic strategies for HPV-associated disease.

### Conclusions

In this study, we established a durable organotypic epithelial raft culture that incorporates HPV-transformed cells and primary human keratinocytes, based on a scaffold-reinforced dermal equivalent. We were able to show that our model is suited for preclinical treatment experiments. We observed plausible time- and dose-dependent effects of cisplatin, 5-FU, and sinecatechins treatment on keratinocytes and HPV-transformed cells. Our model was robust enough to physically tolerate the daily application of two distinct topical formulations to their air-exposed surface, indicating its suitability for topical treatment experiments. We were also able to demonstrate that a differential observation of treatment effects on primary keratinocytes and HPV-transformed cells is possible. The treatment effects observed on a qualitative level were reliably quantified using a machine learning-based cell classification approach. The presented model can easily be integrated into the drug development workflow, it has the potential to be expanded to include further cell types, and it may serve as a meaningful link between monolayer cell culture and animal or clinical studies and is therefore a step forward in the research for novel treatment options of HPV-associated disease.

## Acknowledgements

Richard M. Köhler was supported through an MD thesis grant by the *Heinrich F.C. Behr* foundation.

## Methods

### Ethics approval and consent to Participate

The use of human skin tissue was approved by the Ethics Committee of the Medical Faculty Mannheim (2009-350N-MA). Patients were informed and gave consent to the use of their tissue for scientific research.

### Cell culture

The use of human skin tissue was approved by the Ethics Committee of the Medical Faculty Mannheim (2009-350N-MA). Patients were informed and gave consent to the use of their tissue for scientific research. Primary fibroblasts isolated from explant cultures of de-epidermized dermis were expanded up to passage 15 in Gibco Dulbecco’s Modified Eagle Medium (DMEM) with 10% fetal bovine serum (FBS) and 1% Penicillin/Streptomycin (all Thermo Fisher Scientific Gibco, Schwerte, Germany). For keratinocyte cultivation, fibroblasts were gamma-irradiated with 70 Gy and 10^6^ cells were plated on a 150 mm diameter petri dish (Corning Inc., Corning, USA) to form a feeder layer. To obtain primary keratinocytes, adult human skin tissue was treated with thermolysin to isolate the epidermis which was then incubated with trypsin to release the cells^12^. Keratinocytes were seeded onto the feeder fibroblasts at a count of 250 x 10^3^ cells per culture and cultivated in FAD medium, a 1+3 mixture of Ham’s F12 and DMEM, supplemented with 5% FBS, 1% Penicillin/Streptomycin (all Thermo Fisher Scientific Gibco, Schwerte, Germany), 10^-10^ M cholera toxin, 1.8 x 10^4^ M adenine, 5 µg per ml insulin, 0.4 µg per ml hydrocortisone (all Sigma-Aldrich Co, Steinheim, Germany), and 1 µg human recombinant EGF (PromoCell, Heidelberg, Germany). A selective rho-kinase inhibitor (Y-27632 dihydrochloride; abcam plc, Cambridge, UK) was added at a concentration of 10 µM to prevent premature epithelial differentiation. The authenticity of cell lines CaSki, SW756, SiHa, and HeLa was confirmed by Multiplex human cell line authentication tests (Multiplexion GmbH, Friedrichshafen, Germany). All cervical cancer cell lines were cultured in DMEM supplemented with 10% FBS and 1% Penicillin/Streptomycin during their cultivation in monolayer culture, i.e. prior to their seeding onto the dermal equivalents for generation of OTCs.

### Dermal equivalent generation

Scaffold-enforced dermal equivalents were prepared based on a modified protocol of a previously described method which permits long-term cultivation of skin equivalents.^13^ Two single-layer Bemcot M-3 industrial wipes (Asahi Kasei, Düsseldorf, Germany) were stacked to form a double layer sheet and circular scaffolds with a diameter of 11 mm were punched out from these sheets. A fibrin gel with homogeneous distribution of non-irradiated primary fibroblasts was prepared using TISSEEL Kit (Baxter International Inc., Deerfield, Illinois, USA). Thrombin S (Baxter International) was diluted to 10 U with phosphate-buffered saline (PBS; SERVA Electrophoresis GmbH, Heidelberg, Germany), fibroblasts were resuspended in FBS (Fisher Scientific) to reach a concentration of 1.25 x 10^6^ cells/ml and the solutions were then mixed in equal parts. Bovine fibrinogen (F8630, Sigma-Aldrich Co, Steinheim, Germany) was diluted to a concentration of 8 mg/ml with PBS. Millipore inserts (ThinCert, 12-well, pore size 0.4 µm, translucent; Greiner Bio-One GmbH, Frickenhausen, Germany) were placed in deep-well plates (12-well ThinCert Plate; Greiner Bio-One) and scaffolds were positioned in the inserts. 200 µl of the thrombin-fibroblast suspension were pipetted onto the scaffolds. The same volume of fibrinogen solution was added immediately and the two were mixed quickly through repeated aspiration and pipetting. The cultures were then incubated at 37°C for 30 to 60 minutes to allow formation of fibrin gels. The gels were then submerged in 5 ml DMEM supplemented with 10% FBS, 1% Penicillin/Streptomycin, 1 ng per ml TGFß1 (Sigma-Aldrich Co, Steinheim, Germany) and 50 µg per ml ascorbic acid (Sigma-Aldrich Co, Steinheim, Germany). One day before keratinocyte and tumor cell seeding, custom-made rings (generated from polytetrafluoroethylene (PTFE) tubes) fitted to the inserts were placed directly onto the dermal equivalents to standardize the cell proliferation area and restrict overgrowth on the sides of the dermal equivalent. Additionally, the medium was shifted to rFAD, a 1+3 mixture of Ham’s F12 and DMEM, supplemented with 10% FBS, 1% Penicillin/Streptomycin (all Thermo Fisher Scientific Gibco, Schwerte, Germany), 10^-10^ M cholera toxin and 0.4 µg per ml hydrocortisone (all Sigma-Aldrich Co, Steinheim, Germany). rFAD was additionally supplemented with 50 µg per ml ascorbic acid and 500 E per ml aprotinin (both Sigma-Aldrich Co, Steinheim, Germany).

### Organotypic co-culture generation and harvest

Target keratinocyte-tumor cell mixtures were seeded on the dermal equivalents at a count of 250 x 10^3^ cells per culture and the submerged cultures were incubated for 24 hours. PTFE rings were then removed, and the fluid level was lowered below the OTCs’ upper surface to expose the epithelial cells to air. Aprotinin was reduced to 250 U per ml, and the medium was changed every second day. Cultures were incubated with 10 μM 5-ethynyl-2’-deoxyuridine (EdU) added to the culture medium 4 hours prior to fixation. OTCs were harvested by carefully cutting the Millipore membrane of the insert on its inner rim with a disposable scalpel (Feather Safety Razor Co., Osaka, Japan). OTCs were then fixed for 48 hours in formalin solution (“Spezialfixiermittel fuer Anatomie & Histologie”; Morphisto, Frankfurt am Main, Germany), dehydrated, and embedded in paraffin.

### Cisplatin, 5-FU and sinecatechins treatment

A suspension of 1% SiHa cells and 99% keratinocytes was seeded at a density of 2.2 x 10^5^ per cm^2^ onto previously prepared dermal equivalents as described above. Before the cisplatin treatment experiment, OTCs were cultured for 11 days after start of the air-exposed cultivation. OTCs were then subjected to treatment with either 1, 5 or 10 µM cisplatin applied into the cell culture medium or were left untreated as controls. OTCs were harvested after either 1 week of treatment, 2 weeks of treatment or 2 weeks of treatment plus 2 weeks of culture without further treatment (recovery). For each treatment condition and culture duration, two biological replicates (i.e., two OTCs) were generated.

Before treatment with a commercially available 5% 5-fluorouracil (5-FU cream (Efudix, Viatris Healthcare GmbH, Bad Homburg), OTCs were cultured for 11 days after start of the air-exposed cultivation. OTCs were then subjected to either 10 mg of 5-FU cream applied topically onto the air-exposed epithelial OTC surface or were left untreated as controls. OTCs were harvested after either 1 week or 2 weeks of treatment. For each treatment condition and culture duration, two biological replicates were generated, except for the one-week treatment group which initially contained three replicates. However, only two replicates could be included in further analysis as a result of a processing failure during embedding of the third specimen in paraffin.

Before treatment with sinecatechins using a commercially available 10% sinecatechins ointment (Veregen 10% Salbe, Dr. Pfleger Arzneimittel GmbH, Bamberg, Germany), OTCs were cultured for 12 days after start of the air-exposed cultivation. OTCs were then subjected to either 10−15 mg of ointment applied topically onto the air-exposed epithelial OTC surface or were left untreated as controls. OTCs were harvested after either 1 week or 2 weeks of treatment. Two biological replicates were generated for each harvest time point for treated cultures, and one biological replicate as untreated control.

Treatment with 5-FU and sinecatechins was conducted daily on 5 days per week (see Figure 3). To enable and standardize topical treatment onto the air-exposed epithelial OTC surface, small patches of wax-coated flexible carrier films (“Parafilm M”, Amcor) with a diameter of 11 mm were punched out and irradiated with 300 Gy for sterilization purposes. For application of 5-FU cream and sinecatechins ointment, the respective compound was spread onto the carrier platelets with a spatula. Excess product above the target amount (10 mg) was removed and the carrier platelets were then placed onto the OTCs’ air-exposed surface. After an exposure time of 2 hours in an incubator at 37°C with 5% CO_2_, the carrier patches were removed from the OTCs with a forceps. Potential residues of the topical agent on the OTC surface were then carefully rinsed off by pipetting with PBS. OTCs in the cisplatin group were treated once per week (see Figure 3). Immediately before treatment, cisplatin was suspended in aliquots of 100 µl medium each and added to the growth medium of the OTCs on the first day of each week of treatment.

### Hematoxylin and eosin staining

Sections at a thickness of 4 µm were chemically deparaffinized through a series of xylene clearing reagents, then through a series of alcohol solutions in decreasing concentrations (100%, 96%, 80%, 70%) to water. Sections were then stained with Mayer’s hematoxylin (Dako North America, Inc., Carpinteria, CA, USA), and subsequently blued in a staining box placed under running tap water for five minutes. The sections were then washed in distilled water for three minutes and subsequently stained with eosin (Sigma-Aldrich Co, Steinheim, Germany), followed by dehydration through a series of alcohol solutions in increasing concentrations (70%, 80%, 96%, 100%), then a series of xylene baths, and finally mounting with Eukitt (SP Filling GmbH, Bobingen). Slides were then scanned and digitized with Nanozoomer S210 and examined using NDP.view 2 imaging software (both Hamamatsu Photonics, Hamamatsu, Japan).

### Immunofluorescence

Sections of formalin-fixed paraffin-embedded specimens were generated at a thickness of 3 µm, placed onto coated microscope slides (SuperFrost Plus, Thermo Fisher Scientific Gibco, Schwerte, Germany) and dried overnight at 37°C. Heat induced antigen retrieval was performed by boiling in 100 mM Tris buffer pH 9 for 15 minutes. Sections were first rinsed with double-distilled water and then pre-blocked with 3% bovine serum albumin (Jackson ImmunoResearch, Cambridgeshire, UK) in PBS (SERVA Electrophoresis GmbH, Heidelberg, Germany) for one hour in a humid chamber. The blocking buffer was then removed, and sections were incubated with primary antibodies in a humid chamber for 1 hour at 37°C, then for 18 hours at 4°C in a humid chamber. The primary antibody dilution was subsequently aspirated, and sections were washed three times for 5 minutes each in PBS. Following primary antibodies were used in the given dilutions: rabbit polyclonal anti Collagen Type IV, 1:300 (Rockland Immunochemicals, Pottstown, USA), guinea pig polyclonal anti Keratin 7, 1:100, and mouse monoclonal anti K8, 1:20 (both PROGEN Biotechnik, Heidelberg, Germany), rabbit polyclonal anti K14, 1:1000 (BioLegend Europe B.V., Amsterdam, Netherlands), rabbit polyclonal anti Ki67, 1:200 (abcam plc, Cambridge, UK), rabbit polyclonal anti cleaved caspase 3, 1:200 (Cell Signaling Technology Europe B.V., Leiden, Netherlands), and mouse monoclonal anti p16 undiluted (CINtec/Roche, Mannheim, Germany). For detection, species-specific, fluorochrome-conjugated secondary antibodies (F(ab’)_2_--fragments of donkey-anti rabbit, anti-mouse and anti-guinea pig-antibodies, respectively, conjugated with Cy3 or Alexa Fluor 488 (Jackson Laboratories distributed by Dianova, Hamburg, Germany) were applied in dilutions of 1:800 to 1:1000 together with 4′,6-diamidino-2-phenylindole (DAPI) (2 µg/ml) for nuclear staining. Sections were incubated for 30 minutes at 37°C in a humid chamber followed by 30 minutes at room temperature. The sections were then washed three times with PBS for 5 minutes each, followed by a brief rinse with double distilled water. When DNA-synthesis was to be visualized the Click-iT EdU Cell Proliferation Kit (Thermo Fisher Scientific, Marietta, OH, USA) was applied according to the manual before the antibody staining was performed. The slides were mounted in Dako Fluorescence mounting medium (Dako North America, Inc., Carpinteria, CA, USA) and stored at 4°C in the dark until examination. Slides were examined using Olympus AX70 fluorescence microscope and cell^F imaging software (Olympus GmbH, Hamburg, Germany).

### Object classifier training

For quantification of treatment effects, four sections were obtained from every specimen in the cisplatin, 5-FU, and sinecatechins treatment experiments with 100 µM distance between each section (N=38 specimens). Sections were stained with hematoxylin and eosin (H&E), scanned, and digitized as described above. One section from each specimen of the cisplatin experiment was used for classifier training and validation, and three sections were set aside for final treatment quantification (Figure 5A).

Two researchers (A: ESP, B: RMK) manually annotated one section per specimen used for classifier training in the open source software QuPath^14^ independently from each other. Small groups of epithelial cells were encircled and were marked as groups of SiHa cells, keratinocytes, or degraded cells. Within these regions, objects were automatically detected from star-convex polygon features using a pre-trained convolutional neural network (“2D_versatile_he”) in StarDist.^15,16^ These objects corresponded to one of three classes annotated by the researchers: i) SiHa nuclei, ii) keratinocyte cell nuclei, or iii) degraded cell material regions. Features derived from object morphology and hematoxylin staining were calculated for each detected object in QuPath (e.g., area, length, circularity, mean, and standard deviation of hematoxylin intensity). Extreme gradient boosting tree models (XGBoost^17^) were then trained with the extracted features to separate the three classes of objects (SiHa cells, keratinocytes, degraded cells). Hyperparameters were optimized with Bayesian optimization. To avoid overfitting and thus an inaccurately high classification performance attributable to feature similarity within each specimen, cross-validation was performed in a leave-one-group-out approach where each specimen represented one group. Specifically, to calculate classification performance for a single left-out specimen, a single model was trained using features from all other specimens. The model then classified the objects in the left-out specimen and balanced accuracy was calculated. This procedure was repeated for each single specimen. Classification performances between researcher A and B were statistically compared with a two-sided permutation test with 5 x 10^5^ permutations. A single final model for treatment quantification was then trained using annotations by researcher A from all training slides.

### Treatment effect quantification

Three out of four H&E-stained sections from each specimen (i.e. from 2 specimen/biological replicates per treatment and harvest condition, or 1 specimen for control cultures of the sinecatechins experiment, as described above) had been set aside before object classifier training for the final quantification of treatment effects. In these sections, the epithelium including tumor cells − or the remaining dispersed cell groups in the case of pronounced cell-degrading treatment effects − was annotated and labeled as region of interest (ROI; Figure 5A). Within the ROI, objects were detected using StarDist and features extracted using QuPath (as described above). Objects were then classified as either SiHa cells, keratinocytes or degraded cells using the final model.

SiHa and keratinocyte cell counts of each slide were divided by the ROI’s total area to account for difference of ROI area between different slides and specimens, to derive cell counts per mm². SiHa and keratinocyte cell counts of *treated* cultures were then normalized through division by the respective mean cell count of the *untreated control* cultures of the respective experiment. A value larger than 1 thus indicates an increased cell count relative to the untreated control, whereas a value smaller than 1 indicates a decreased cell count. Finally, we calculated the ratio of SiHa to keratinocyte by division of SiHa count by keratinocyte count. This metric was derived in order to approximate the relative strength of the treatment effect on transformed cells versus healthy tissue. A value larger than 1 indicates that more SiHa cells were present, a value smaller than 1 indicates more keratinocytes. For treatment quantification, no statistical analysis was performed as the experiments were intended as a proof of concept to demonstrate the feasibility of semi-automated quantification of treatment effects, and sample size was therefore low (N=1/2 specimens, i.e. biological replicates) per treatment and harvest condition.

### Data and code availability

Code and data that support the findings of this study are openly available at https://github.com/richardkoehler/paper-otcs.

