## Supplements for "An organotypic in vitro model of human papillomavirus-associated precancerous lesions allowing automated cell quantification for preclinical drug testing"

### Supplementary Information: Figures S1-S3

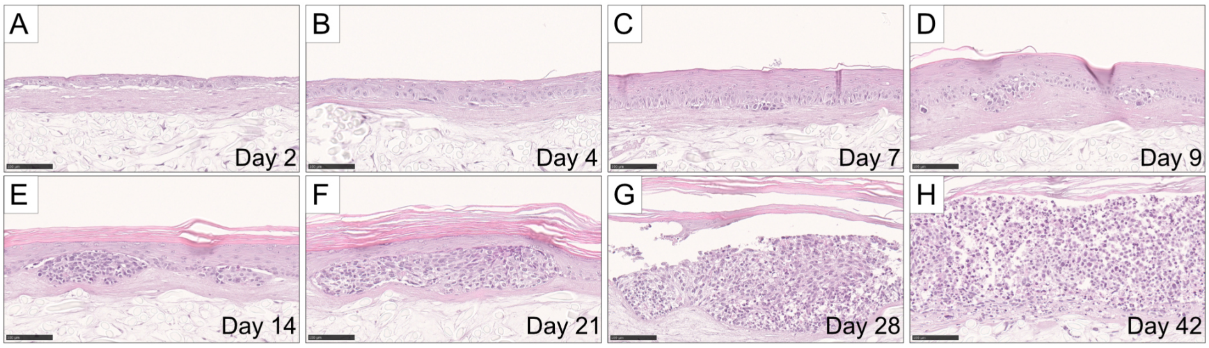


**Figure S1: Morphological evolution of organotypic epithelial raft cultures (OTCs) mimicking HPV-induced precancerous lesions over the course of 6 weeks.** Exemplary depiction of the tissue architecture within established OTCs at a seeding ratio of 99% primary keratinocytes and 1% SiHa cells. (A) Two days after epithelial cell seeding, a confluent layer of cells covered the dermal equivalent. (B, C, D) Keratinocytes continued stratification until all layers of a squamous epithelium had been formed around day nine. (E) After about two weeks of cultivation, readily discernible nests of SiHa cells were observed. (F, G) After that, SiHa cells continued to expand and after about four weeks, they had covered almost the entirety of the dermal equivalent. (H) After six weeks of culture, intact cells continued to be observable in the OTC. Scale bars: 100 µm.

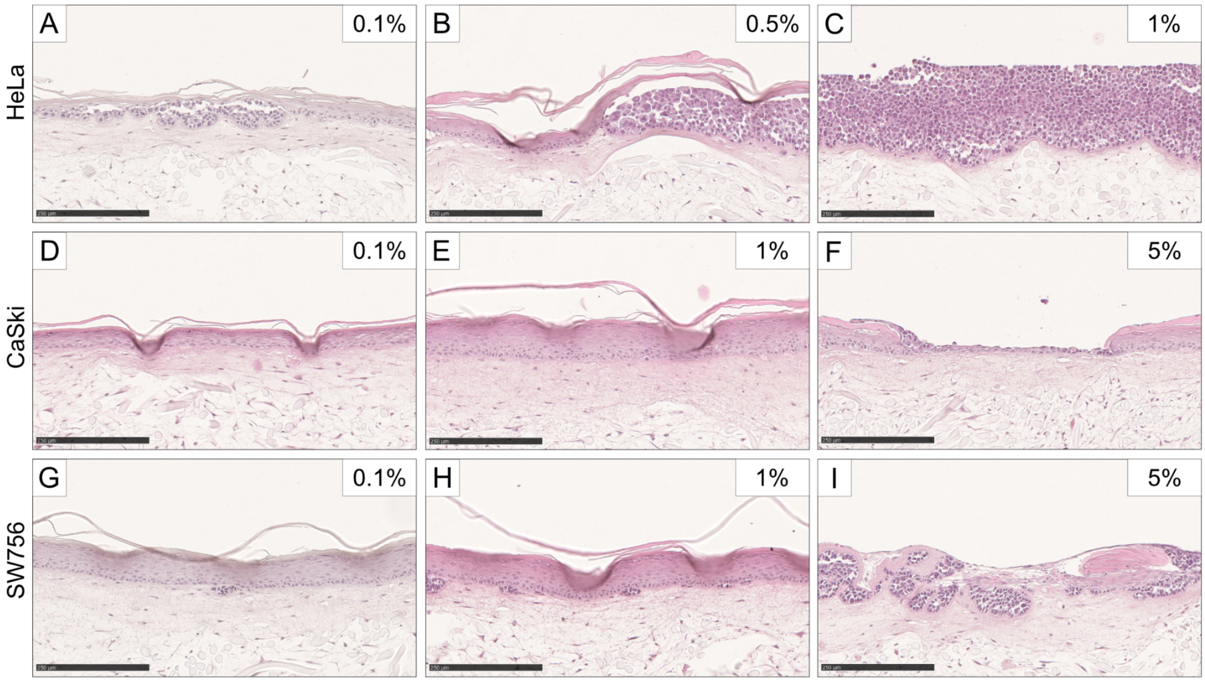


**Figure S2: Establishment of OTCs mimicking HPV-induced precancerous lesions from different cell lines.** Keratinocytes co-seeded with cells from different tumor cell lines and in varying ratios. OTCs were harvested after 13 days. Cell lines include HeLa 0.1 – 1% (A-C), CaSki 0.1 – 5% (D-F), and SW756 0.1 – 5% (G-I).%: proportion of tumor cells seeded in relation to healthy keratinocytes. Scale bars: 250 µm.

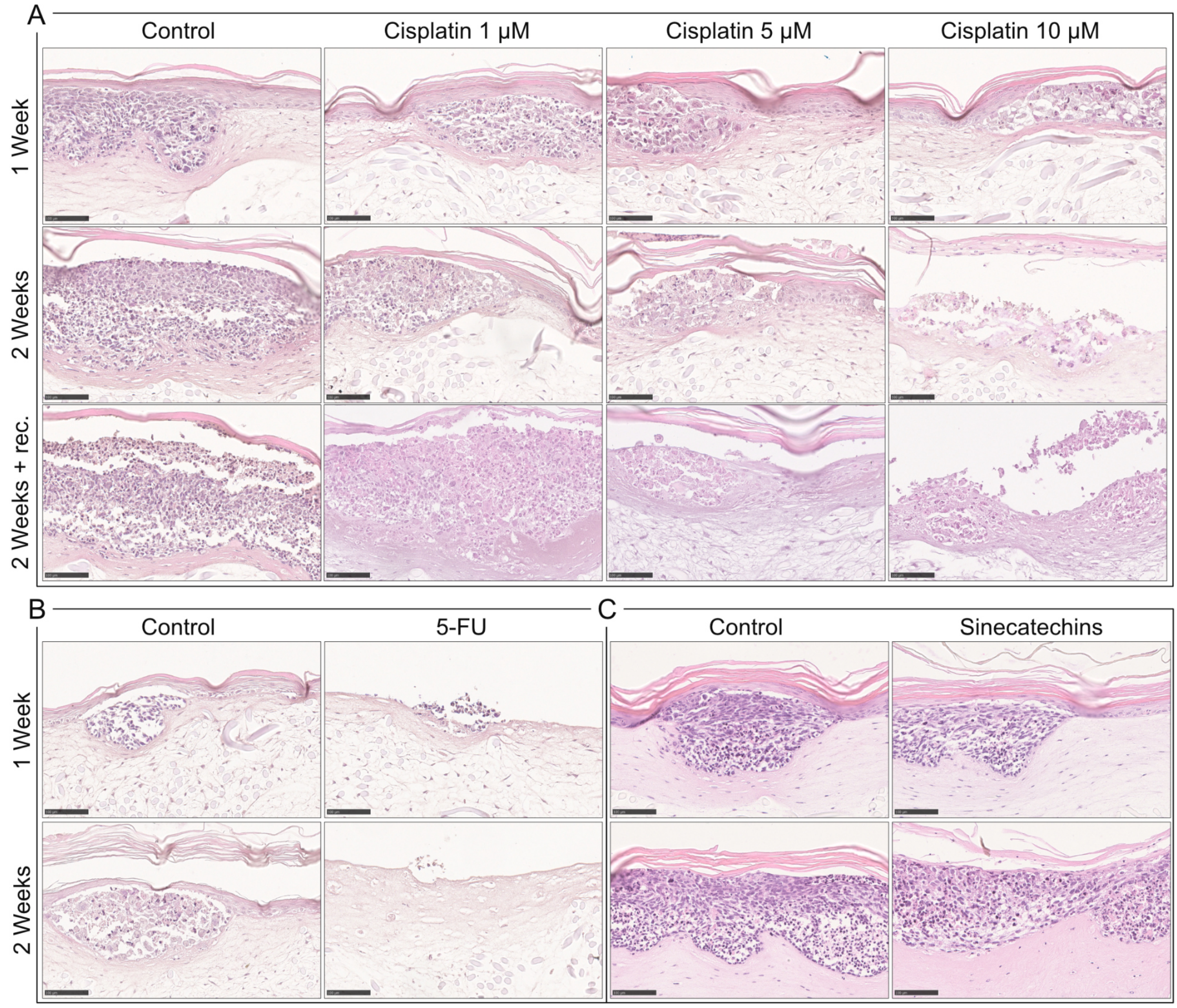


**Figure S3: Full overview of treatment effects on OTCs mimicking HPV-induced precancerous lesions after simulation of topical and systemic drug application.** Organotypic epithelial raft cultures (OTCs) with epithelial cells seeded at a ratio of 99% keratinocytes and 1% SiHa cells treated with cisplatin added to the growth medium, or with topical application of either a commercially available 5-fluorouracil (5-FU) cream or sinecatechins ointment. Controls were left untreated. Cultures were harvested after one week of treatment, two weeks of treatment, or two weeks of treatment plus two weeks without further treatment (recovery phase). (A) Cisplatin addition to the growth medium led to a time- and dose-dependent cell degradation. (B) Daily administration of 5-FU ointment resulted in a complete detachment of the epithelium. (C) Topical application of sinecatechins led to a minor degradation of SiHa cells compared to untreated controls. Scale bars: 100 µm. Note: A subset of images is also shown in Figure 3 of the main text.
